# Spatial distribution and seasonal fluctuations of *Drosophila* on Santa Catalina Island with an emphasis on the repleta species group

**DOI:** 10.1101/007732

**Authors:** JY Kao, SV Nuzhdin

## Abstract

Santa Catalina Island is a small island off the coast of southern California and for its modest size harbors several species of flies from the *Drosophila* genus. We performed an island-wide survey of *Drosophila* species to ascertain which species were endemic to the island and where they were most abundant. In doing so, we have assembled useful sampling information for researchers who wish to conduct field studies on Santa Catalina Island. From this survey, we determined that *Drosophila hamatofila, Drosophila mainlandi,* and *Drosophila mettleri* were the prominent repleta species on the island. Other repleta species encountered included *Drosophila mojavensis* and *Drosophila wheeleri.* Non-repleta species sighted on the island include *Drosophila melanogaster, Drosophila pseudoobscura, Drosophila simulans,* and an unknown species not seen before on the island. Additionally, we performed seasonal collections at two locations on the island and observed that species abundance and composition at these two sites vary between seasons. One of the seasonal sites was sampled in two consecutive summer seasons, which revealed that species composition had shifted between years, but relative species abundances were approximately the same.

## Introduction

Santa Catalina Island is part of the Channel Islands located off of the southern coast of the U.S. state of California about 35 kilometers south-southwest from the city of Los Angeles, CA, USA. The island is approximately 35.4 kilometers long and 12.9 kilometers across at the widest point and has two towns on either end of the island: Two Harbors, on the north side, and Avalon, on the south side. The climate on the island is classified as subtropical with mild winters and warm temperatures all year round. The proximity to several research institutions and research facilities on the island provided by the Wrigley Marine Science Center and the Catalina Island Conservancy adds to the convenience of field research on the island. Both marine and terrestrial researches have been conducted on the island with an emphasis on the marine front. The vertebrate and plant species on the island are well documented, but with regards to invertebrates, the documentation is sparse. Some studies have focused on specific arthropods endemic to the island, but there have been no island-wide surveys of arthropod species on the island. Out of the arthropods on the island, one of the most studied is the *Drosophila* flies [Reed et al. 2008; Hurtado et al. 2004] and in particular the repleta species group, which utilize cacti as their plant host. There is reason to believe that there are endemic populations of *Drosophila* species inhabiting the island as evidenced by adaptation to island-specific host cacti in certain species [Matzkin 2014; Castrezana and Bono 2012]. Though having previously been studied, the spatial and temporal aspects of this species group (as well as other *Drosophila* species) has not been investigated on the island. Santa Catalina Island in previous surveys was mostly considered as one location to determine species distribution over a larger geographical area [Heed 1982].

Previously sampled species available at the Drosophila Species Stock Center (San Diego, CA) indicate that at eight different *Drosophila* species live on the island. These species include *D. melanogaster, D. simulans, D. hamatofila, D. mojanvensis, D. mettleri, D. mainlandi, D. pseudoobscura,* and *D. wheeleri.* However, the spatial distribution of these species across the island is not known since the collection information available does not include coordinates, but lists the entirety of Santa Catalina Island as a single sample location. From an informal island-wide insect survey conducted April 2011, we have data suggesting that species distribution on the island is not ubiquitous despite the prevalence of host cacti on the island. We returned to the island for a more formal and deeper island-wide survey of *Drosophila* species in June and July of 2012. We noted not only the coordinate locations of collection sites, but also the elevation because altitudinal factors also play a role in population density [Guruprasad et al. 2010]. We have also conducted seasonal collections at two select locations on the island to roughly estimate species composition over a year because seasonality can also plays a role in density of populations [Guruprasad et al. 2010; Torres and Madi-Ravazzi 2006, Dobzhansky and Pavan, 1950, Patterson 1943]. From our survey, we aim to illuminate the spatial distribution and seasonal patterns of select *Drosophila* species to aid the planning of specimen collection efforts for field researchers.

## Materials and Methods

### Ethics Statement

With a collection permit from the Catalina Island Conservancy, we were able to collect on Conservancy land. We also contacted the Santa Catalina Island Company to collect on their land. The Wrigley Marine Science Center also gave us permission to collect at the Center. None of the species sampled were protected.

### Island-wide collections

A preliminary collection was conducted in April 2011 to initially assess species and ease of trapping on Santa Catalina Island. Fly traps were assembled from plastic water and soda bottles with various types of bait (i.e. banana, watermelon, honeydew, papaya, strawberries). Traps were placed in trees or bushes near potential food sources (i.e. cactus, fruit trees, trashcans, etc.) and left for 24 hours before retrieval. Female flies were aspirated from the traps and isofemale lines were established on banana-opuntia food (recipe available online from the Drosophila Species Stock Center, San Diego, CA).

A year later in the months of July 2012 and August 2012, we assembled fly traps made from 480mL (16.9 fl. oz.) plastic water and soda bottles baited approximately 3 cm deep with a mix of rotten banana mash, yeast, and *Opuntia* cactus powder. Sticks were added to the trap for perch sites and three holes were cut into the side of the bottles for entry [Markow & O’Grady 2006]. This particular bait was determined in preliminary trappings to be the most effective in attracting all of the previously known species to inhabit the island (list from Drosophila Species Stock Center, San Diego, CA) compared to other baits that were tried (i.e. opuntia cactus + yeast, banana + yeast, watermelon, papaya, strawberry, honeydew, etc.). Four traps were distributed per site at 24 sites across Catalina Island (Figure 1). Traps were hung on trees branches or in bushes at a minimum of one meter above ground to prevent scavengers such as the island fox from damaging our collection efforts. The minimum distance between traps ranged from six meters to 300 meters. We chose placement based on the immediate proximity of potential food sources (i.e. cactus patches, trashcans, fruit trees, etc.) to draw out flies living in these natural substrates. After 24 hours of trap deployment, the traps were subsequently collected. Traps that contained zero flies after the initial 24 hour period near the two towns Avalon and Two Harbors were left out for 48 hours as well as redeployed in different locations on two (Avalon) to three (Two Harbors) separate excursions. Only one collection effort per site at Avalon and Two Harbors were included in this dataset. Flies were removed from the traps via aspirators and sorted under a scope into repleta species group and non-repleta species subgroup.

**FIGURE 1.**
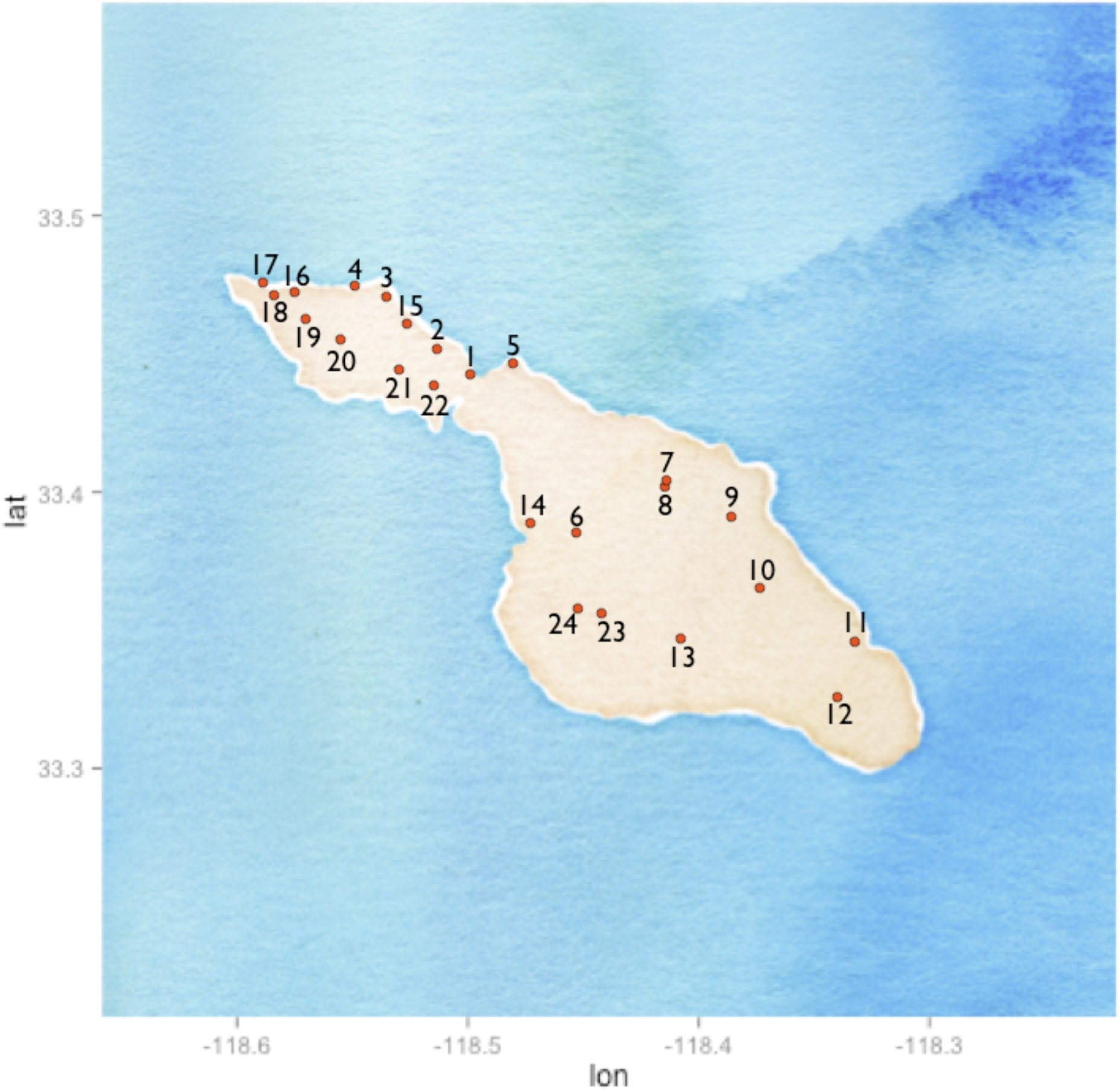
Map of collection sites on Santa Catalina Island.

### Seasonal collections

In addition to summer collections, we returned to our sites at WMSC and Little Harbor campgrounds in November 2012, January 2013, and April 2013 for fall, winter, and spring collections. Flies were also collected for an additional summer season in July 2013 at WMSC. Traps were assembled, deployed, and retrieved as previously described.

### Species identification

In our preliminary collections from April 2011, established isofemale lines that were determined to be of the repleta species group (as indicated by spotted thoraces), were sent to the Drosophila Species Stock Center (San Diego, CA) for species identification.

The species in the repleta group are nontrivial to distinguish because pigmentation patterns are very similar between these species. Only males can be visually identified by looking at the aedeagus morphology via genitalia dissections. Females of this subgroup are not distinguishable from each other except by establishing isofemale lines and examining male progeny aedeagus morphology. Collected males from the island-wide collections in July/August 2012, November 2012, January 2013, and April 2013 were dissected for species identification and females were preserved in ethanol and are available for sequencing. A set of females was also identified from the July/August 2012 collection by establishing isofemale lines and examining male progeny.

Non-repleta species are fairly distinct in pigmentation and morphology and were examined under a scope for identification, but numbers were not recorded, only presence or absence on the island was taken into account.

### Collection analysis

We classified collection sites as “hot”, “warm”, or “cold” according to the number of flies present in traps. Hot spots are defined as locations where the two traps collected more than 20 male flies. Warm spots are defined as locations where the traps contained 10 to 20 male flies. Cold spots are sites where there were less than five flies were present in the traps set. All analysis was performed in R. To determine differences of species compositions between sites, we used Chi-squared tests. Pearson’s r was calculated to assess the relationship between altitude and number of specimens caught and corresponding p-values were also calculated to assess statistical significance.

## Results

### Density and distribution of repleta species

The preliminary survey in April 2011 resulted in the establishment of nine repleta species isofemale lines. Out of the nine isofemale lines, seven were identified as *D. mojavensis* and two were found to be *D. hamatofila* (Table 1).

**TABLE 1:** Data from preliminary collections in April 2011.

| Line/Info | Location | Latitude | Longitude | Flytrap medium | Identification |
| --- | --- | --- | --- | --- | --- |
| **1LHBR** | Little harbor campground | 33.38823 | -118.5363 | Banana | mojavensis |
| **2LHBR** | Little harbor campground | 33.38823 | -118.5363 | Banana | mojavensis |
| **5LHBR** | Little harbor campground | 33.38823 | -118.5363 | Banana | mojavensis |
| **6LHBR** | Little harbor campground | 33.38823 | -118.5363 | Banana | mojavensis |
| **7LHBR** | Little harbor campground | 33.38823 | -118.5363 | Banana | mojavensis |
| **8LHBR** | Little harbor campground | 33.38823 | -118.5363 | Banana | mojavensis |
| **9LHBR** | Little harbor campground | 33.38823 | -118.5363 | Banana | mojavensis |
| **1AGBR** | Avalon gate | 33.34744 | -118.3346 | Banana | hamatofila |
| **3WWR** | Wrigley Institute | 33.444908 | -118.4822 | Watermelon | hamatofila |
| **1WBP** | Wrigley Institute | 33.444908 | -118.4822 | Banana | pseudoobscura |
| **1WHP** | Wrigley Institute | 33.444908 | -118.4822 | Honeydew | pseudoobscura |
| **2WHP** | Wrigley Institute | 33.444908 | -118.4822 | Honeydew | pseudoobscura |
| **1AGPP** | Avalon gate | 33.34744 | -118.3346 | Papaya | pseudoobscura |
| **2WWP** | Wrigley Institute | 33.444908 | -118.4822 | Watermelon | pseudoobscura |
| **1JPP** | Road Junction | 33.36588 | -118.3742 | Papaya | pseudoobscura |
| **2JPP** | Road Junction | 33.36588 | -118.3742 | Papaya | pseudoobscura |
| **1EHP** | Emerald Bay | 33.46889 | -118.5363 | Honeydew | pseudoobscura |
| **1ESP** | Emerald Bay | 33.46889 | -118.5363 | Strawberry | pseudoobscura |
| **2ESP** | Emerald Bay | 33.46889 | -118.5363 | Strawberry | pseudoobscura |

From the first summer season (2012), we dissected 279 males and preserved 177 females in ethanol. Another 120 females identified after the establishment of isofemale lines. Species compositions at each of the sites were found to be statistically different from each other (Table 2, Chi-squared, p <0.0001). We identified three collection hot spots on the island and two warm spots. The three collection hot spots on the island include Starlight Beach, Wrigley Marine Science Center (WMSC), and Little Harbor Campgrounds. The two warm spots were at Eagle’s Nest Lodge and Middle Ranch Junction. Out of the four repleta species identified, the most abundant were *D. mainlandi, D. mettleri,* and *D. hamatofila.* Only four *D. mojavensis* males were collected.

**TABLE 2:**
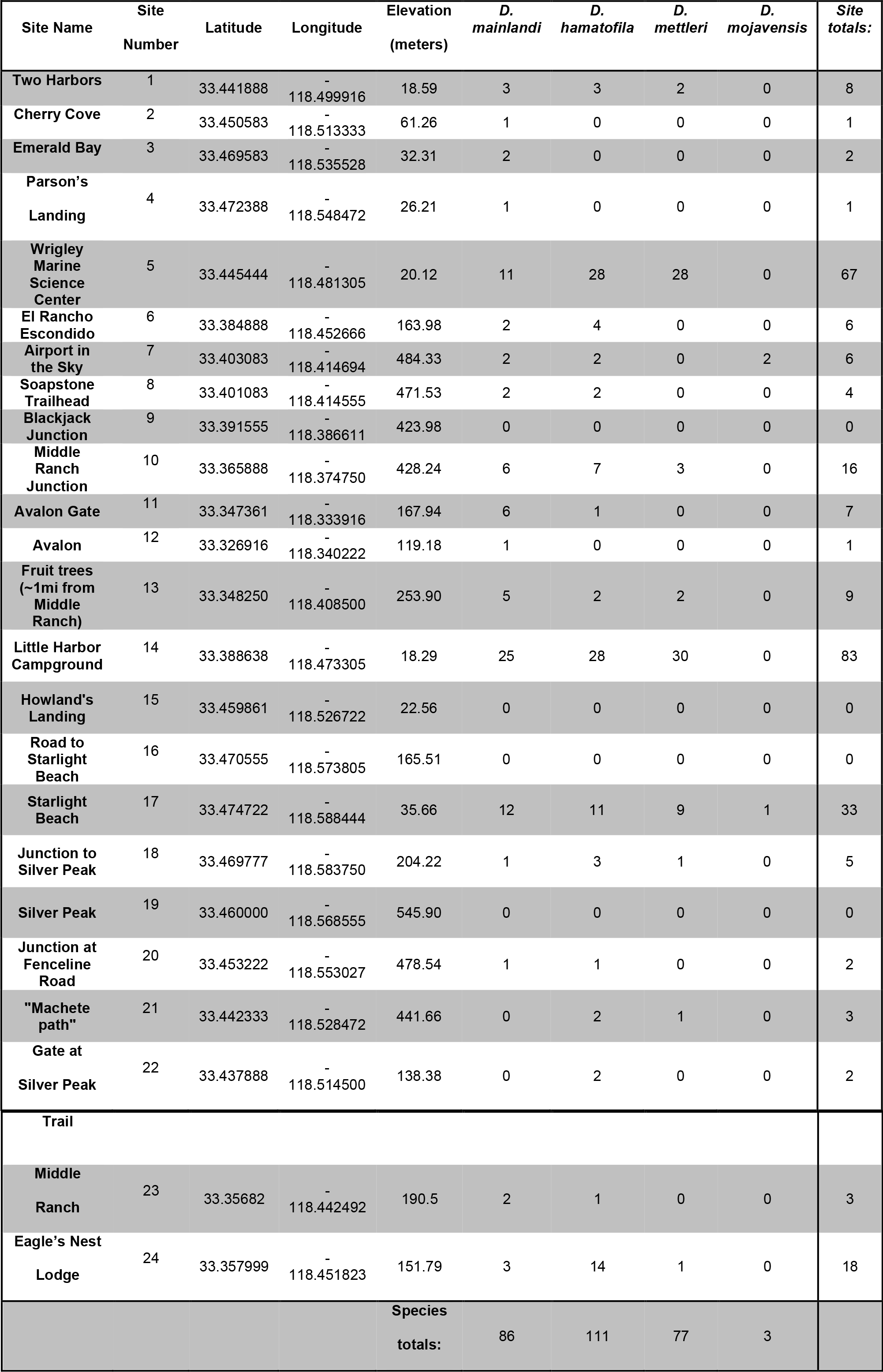
Species compositions at each sampling site from summer 2012 collections.

The species compositions at the collection hot and warm spots were compared. Sites with less than 10 flies were excluded from the comparison. Species compositions at different sites across the island were varied as shown in Table 2. Little Harbor campgrounds and Starlight beach were very similar in species composition with *D. mainlandi, D. hamatofila,* and *D. mettleri* in approximately equal proportions. At Eagle’s Nest Lodge, *D. hamatofila* was the main species collected and at Middle Ranch Junction *D. mainlandi* was the most abundant species. *D. hamatofila* and *D. mettleri* were dominant at the WMSC. It appears that there is a significant correlation of *D. hamatofila, D. mettleri,* and *D. mainlandi* occurring together at collection sites. There also appears to be a marginally insignificant negative trend of number of flies collected and elevation (Table 3).

**TABLE 3:** Correlations and associated p-values of *Drosophila* species on Catalina Island. Pearson’s r is in the top half of the table and associated p-values are in the lower half of the table. Significant p-values and associated correlations are highlighted in light grey.

|  | Elevation | *D. mainlandi* | *D. hamatofila* | *D. mettleri* | *D. mojavensis* | Site totals |
| --- | --- | --- | --- | --- | --- | --- |
| Elevation |  | -0.3455197 | -0.33033703 | -0.355793569 | 0.197617260 | -0.35176745 |
| *D. mainlandi* | 0.09819 |  | 0.84554751 | 0.880838365 | 0.090862560 | 0.93227486 |
| *D. hamatofila* | 0.1149 | <0.001 |  | 0.939890235 | 0.013608634 | 0.97127769 |
| *D. mettleri* | 0.08795 | <0.001 | <0.001 |  | -0.007410574 | 0.98068531 |
| *D. mojavensis* | 0.3546 | 0.6728 | 0.9497 | 0.9726 |  | 0.04779099 |
| Site totals | 0.09186 | <0.001 | <0.001 | <0.001 | 0.8245 |  |

For the fall collection time point, we found only one *D. mainlandi* male at WMSC and 12 *D. mainlandi* males at Little Harbor Campgrounds. No flies were caught in the winter and very few *D. mainlandi* males were collected in the spring with four males at Little Harbor Campgrounds and one male at WMSC. Our spring collections were much lower than what was collected in an island-wide preliminary collection in spring of 2011 (Table 1). In the following summer season (2013), we collected a total of 168 male repleta species specimens from the WMSC sites. There were 81 *D. mainland,* 55 *D. mettleri,* 25 *D. hamatofila,* 4 *D. mojavensis,* and 3 *D. wheeleri.* These species compositions were vastly different from the composition in the previous summer (2012) at WMSC (Figure 2).

**FIGURE 2.**
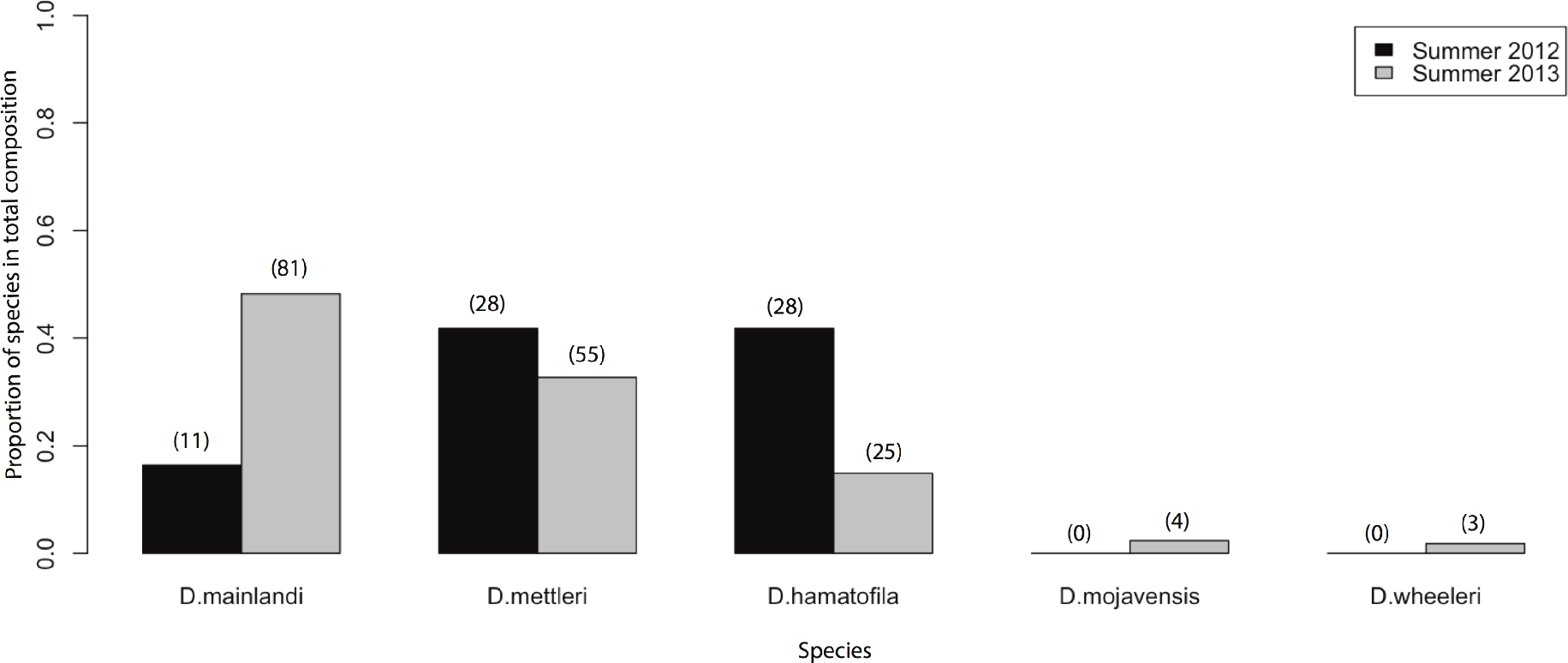
Species composition at the Wrigley Marine Science Center between summers. Absolute numbers of specimens collected are designated in parentheses above each bar

### Non-repleta species

Ten isofemale lines of *D. pseudoobscura* were established from the preliminary April 2011 survey, but no *D. melanogaster* or *D. simulans* were collected in the same survey. In subsequent collections in 2012 and 2013, *Drosophila melanogaster, Drosophila simulans,* and *Drosophila pseudoobscura* were found on the island with the former two species in more relative abundance than the latter between the 2011 preliminary collection and 2012/2013 collections (personal observation).

One unidentified male was collected from one of our sites at Emerald Bay during the summer collections (Figure 3). After consulting with the Drosophila Species Stock Center in San Diego, CA, the specimen had tergite pigmentation most similar to *Drosophila busckii.* However, thorax pigmentation was darker than the species standard. Efforts of collecting more flies like the unidentified specimen in the summer were unsuccessful, but a second male specimen was collected when we returned for collections in the fall at WMSC.

**FIGURE 3.**
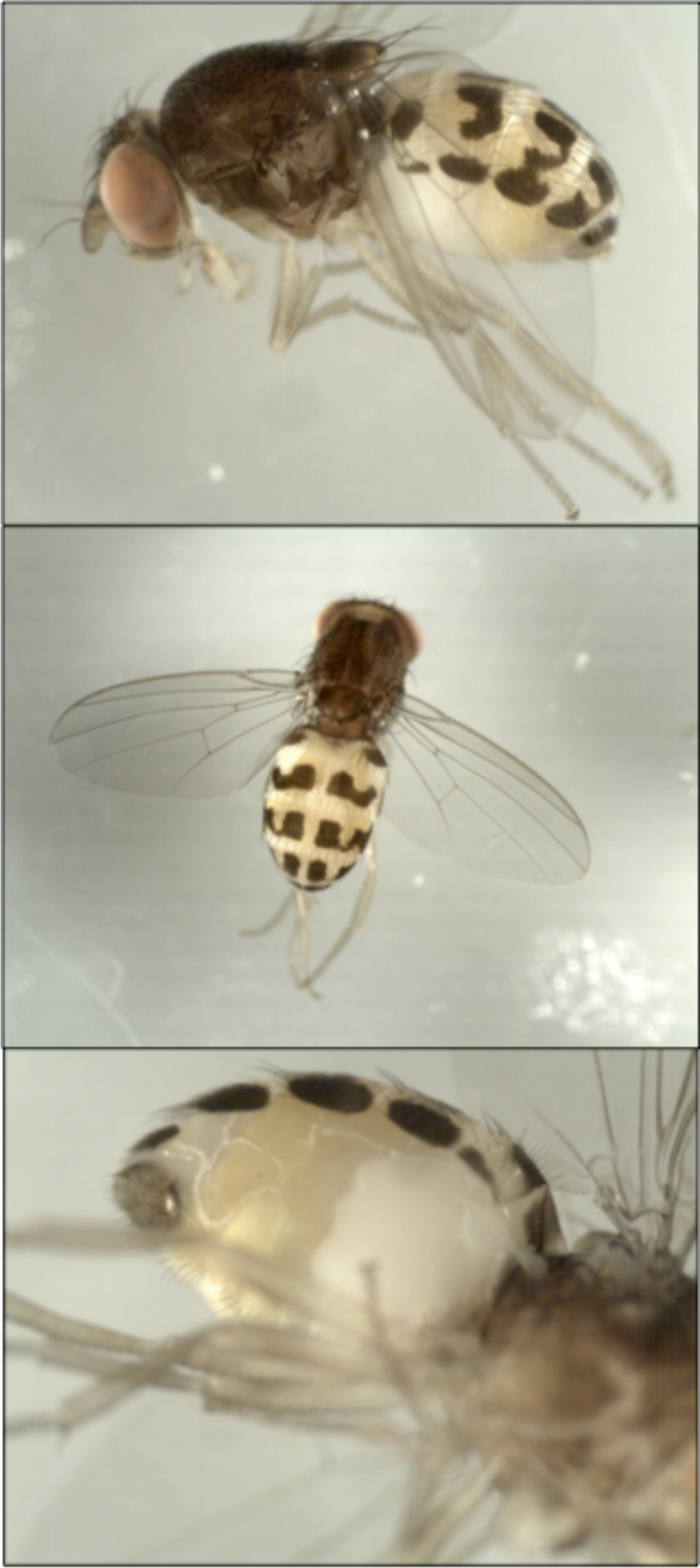
Photos of unknown specimen.

## Discussion

We have assembled information on the distribution of select fly species on Santa Catalina Island for future field researchers interested in collecting specimens from the *Drosophila* repleta species group. We have also sighted *D. melanogaster, D. simulans,* and *D. pseudoobscura* on the island as well as observed a new species not previously seen before on the island.

All of the species we encountered on the island are known human commensals and thus should be abundant in areas with high human occupancy such as towns and cities [Powell 1997]. Therefore, it was surprising to find that the towns on Catalina (i.e. Avalon and Two Harbors) had very low to no yields of fly collections despite multiple sampling attempts for varying lengths of time. Reasons why Avalon and Two Harbors have a low population of flies are unclear at the moment. Further investigation into weather and wind patterns and possible pesticide use might give clues as to why these areas are not inhabited by many flies.

Overall, it appears that there are locations on Santa Catalina Island that may be more conducive to specimen collections than other areas. Furthermore, our findings point towards a seasonality of overall fly population with the summer season possibly being the best for specimen collection in terms of numbers and diversity. The species composition within the same location and season between years appear to be not stable according to our spring (years 2011 and 2012) and summer (years 2012 and 2013) sampling data. The different values between years suggest that species composition between seasons and years is highly sensitive to environmental factors (i.e. rainfall, temperatures, humidity, etc.). More sampling at regular time intervals over several years would be needed to determine if this were the case. In addition to environmental factors, migration to and from the mainland could be an influential factor in overall fly population numbers as well as composition in the summer (i.e. high tourism season) versus winter (i.e. low tourism season). Additional collections over several years accompanied by paired sequencing of island and mainland specimens would be needed to assess the plausibility of migration having an effect on overall island populations. These suggested efforts would also help establish the permanence and identity of the unknown species we encountered on the island.

In summary, our data suggests that the summer season between July and August is the best time to perform specimen collections in terms of population abundance. The site that yielded the most total number of specimens was Little Harbor Campgrounds and the second most was the Wrigley Marine Science Center providing evidence that these two sites would be the good candidate collection locations in terms of maximizing specimens in the field.

## Acknowledgements

The authors would like to thank the Wrigley Marine Science Center for the use of their facilities and equipment on the island as well as the Catalina Island Conservancy for the training and use of a 4WD vehicle to access remote parts of the island. We would also like to thank the lab of Mariana Mateos at Texas A&M University, in particular, Lauryn and Caitlyn Winter, for assistance in field collections and species identification of a set of collected specimens. They would also like to thank Maxi Polihronakis Richmond at the UC San Diego *Drosophila* species stock center for instruction on genitalia dissections as well as consulting on identifying *Drosophila* species. This project was made possible by the Wrigley Institute Summer Fellowship as well as support from the National Science Foundation (NSF 1226895) and the National Institutes of Health (NIH RO1 MH091561).

## References

Reed LK, La Flamme BA, Markow TA. (2008) Genetic Architecture of Hybrid Male Sterility in Drosophila. PLoS One. 3 (8): e 3076.

Hurtado LA, Erez T, Castrezana T, Markow TA. (2004) Contrasting Population Genetic Patterns and Evolutionary Histories Among Sympatric Sonoran Desert Cactophilic Drosophila. Molecular Ecology 13: 1365–1375.

Matzkin LM. (2014) Ecological genomics of host shifts in *Drosophila mojavensis*. In: Landry CR, Aubin-Horth N, editors. Ecological Genomics: Ecology and the Evolution of Genes and Genomes. pp. 233–247.

Castrezana S, Bono JM. (2012) Host Plant Adaptation in *Drosophila mettleri* Populations. PLoS One. DOI: 10.1371/journal.pone.0034008.

Heed WB. (1982) The Origin of Drosophila in the Sonoran Desert. In: Barker JSF, Starmer WT, editors. Ecological Genetics and Evolution: The Cactus-Yeast-Drosophila Model System. pp. 65–80.

Guruprasad BR, Hedge SN, Krishna MS. 2010. Seasonal and altitudinal changes in population density of 20 species of Drosophila in Chamundi hill. Journal of Insect Science. 10:123.

Torres FR, Madi-Ravazzi L. 2006. Seasonal variation in natural populations of *Drosophila* spp. (Diptera) in two woodlands in the State of Sao Paulo, Brazil. Iheringia, Sér. Zool., Porto Alegre, 96(4):437–444

Dobzhansky T, Pavan C. 1950. Local and seasonal variations in relative frequencies of species of *Drosophila* in Brazil. Journal of Animal Ecology. 19(1):1–14.

Patterson JT. 1943. The Drosophilidae of the Southwest. Univ. Texas. Publ. 4313: 7–216.

Markow TA, O’Grady PM. 2006. Drosophila: A guide to species identification and use. London: Academic Press Elsevier. 145 p.

Powell JR. 1997. Progress and Prospects in Evolutionary Biology: The Drosophila Model. New York: Oxford University Press. 10 p.

